# Multidimensional Mechanistic Study of Panax Notoginseng Saponins in the Treatment of Alcohol-Induced Osteonecrosis of the Femoral Head: Integrating Network Pharmacology, Molecular Dynamics Simulation and In Vivo Validation

**DOI:** 10.64898/2026.04.28.721283

**Authors:** Ruokun Bai, Hua Su, Jian Mo, Xiaoyun Zhang, Zhengpeng Li, Xiangshan Chen, Sentao Ye, Xiayu Nie, Shuai Chen, Bing Liang

## Abstract

**Background:** Alcohol-induced osteonecrosis of the femoral head (AIONFH) is an orthopedic disorder from chronic alcohol abuse, characterized by disrupted femoral head blood supply, osteocyte death and structural collapse. Current hip-preserving therapy is unsatisfactory, and most patients eventually require total hip arthroplasty. Panax Notoginseng Saponins (PNS), the core active component of Panax notoginseng, exerts pro-angiogenic and anti-osteocyte apoptosis effects, but its specific therapeutic mechanism remains unclear.

**Objective:** This study used network pharmacology, molecular dynamics simulation and animal experiments to identify PNS’s active components, core targets and key pathways for AIONFH, verify its in vivo efficacy, and provide a scientific basis for clinical application.

**Methods:** PNS active components, their targets and AIONFH-related targets were screened from databases; intersection targets constructed an interaction network, core targets were screened by three machine learning algorithms, with concurrent GO and KEGG analysis. Molecular docking was performed between core targets and PNS components; Gromacs 2022 conducted 100 ns simulation to evaluate complex stability. AIONFH rat models were grouped with 4-week intragastric intervention; pathology, immunofluorescence and PCR were used for detection.

**Results and Discussion:** Network pharmacology identified 127 PNS targets and 18 intersections with 672 AIONFH targets. Six core targets (including FGF2, HSD11B1) were screened; KEGG indicated VEGF pathway as key. Ginsenoside Re bound HSD11B1 with the lowest binding energy (-12.4 kcal/mol), and 100 ns simulation confirmed complex stability. Animal experiments showed PNS improved trabecular structure and regulated osteocyte activity. PNS treats AIONFH via multi-component, multi-target mode, core mechanism being osteocyte apoptosis inhibition.

**Results and Discussion:** Network pharmacology screening identified 127 potential targets of PNS, and 18 potential intersection targets were obtained by overlapping with 672 AIONFH-related targets. Six core targets including FGF2 and HSD11B1 were screened out by machine learning, and KEGG analysis indicated that the VEGF pathway and other pathways were the key signaling pathways for PNS action. Molecular docking showed that Ginsenoside Re had the lowest binding energy with HSD11B1 (-12.4 kcal/mol), and 100 ns molecular dynamics simulation confirmed the stable conformation of this complex. Animal experiments demonstrated that PNS could improve trabecular bone structure and regulate osteocyte activity. In summary, PNS exerts a therapeutic effect on AIONFH through a multi-component, multi-target and multi-pathway mode, with the core mechanism of inhibiting osteocyte apoptosis.

## 1 Introduction

Alcohol-induced osteonecrosis of the femoral head (AIONFH) is an orthopedic disease characterized by the interruption or significant impairment of femoral head blood supply caused by chronic alcohol abuse, which subsequently leads to osteocyte death, femoral head structural collapse and progressive loss of hip joint function. Its clinical manifestations include pain in the hip or inguinal region, and in the advanced stage, limb shortening, abnormal gait and limited hip joint movement may occur **[1-3]**. Epidemiological investigations have shown that AIONFH accounts for 20-40% of non-traumatic osteonecrosis of the femoral head cases, predominantly affecting long-term alcohol abusers aged 30-50 years, with a tendency of younger onset **[4]**. Clinically, statins or anticoagulants are used for hip-preserving therapy of AIONFH, but the efficacy is poor, and most patients still need total hip arthroplasty **[5]**. Therefore, it is urgent to explore novel intervention strategies for AIONFH through experimental research.

Panax Notoginseng Saponins (PNS) are the main active components extracted from the rhizome of Panax notoginseng, a traditional Chinese medicine, and are well-known for their pro-angiogenic, anti-inflammatory and anticoagulant effects. At present, PNS has been widely used in the treatment of cardiovascular and cerebrovascular diseases and bone loss **[6-7]**. In addition, PNS has been proven to effectively improve bone microcirculation and inhibit osteocyte apoptosis **[8]**. In in vitro cell experiments, PNS significantly increases intraosseous capillary density, reduces marrow fat accumulation and decreases osteocyte mortality by activating the PI3K/Akt signaling pathway **[9]**. Studies have shown that PNS can alleviate the degree of femoral head collapse by up-regulating the expression of angiogenic and osteogenic proteins **[10]**. In conclusion, PNS is a potential therapeutic drug for AIONFH.

Despite the multiple pharmacological properties of PNS, including anti-inflammatory, pro-angiogenic and bone repair effects, its mechanism of action in the treatment of AIONFH has not been fully elucidated **[11-15]**. To further explore the pharmacological mechanism of drugs in disease treatment, researchers have increasingly resorted to computational methods such as network pharmacology, molecular docking and multi-dimensional bioinformatics analysis in recent years, so as to reveal the pathogenesis of diseases and drug intervention pathways **[16-19]**.

This study aimed to systematically clarify the potential of PNS in the treatment of AIONFH, identify its key therapeutic targets and explore its underlying molecular mechanisms through network pharmacology, molecular docking and molecular dynamics simulation. In addition, animal experiments on rats were conducted to further verify the therapeutic effect of PNS on AIONFH at the in vivo level and perform an empirical analysis of the relevant pharmacological mechanisms. The results of this study provide a solid theoretical and experimental basis for the further development and application of PNS in the treatment of AIONFH.

## 2 Methods

### 2.1 Network Pharmacology

#### 2.1.1 Acquisition of PNS Active Components and Corresponding Targets

The main active components of PNS and their Canonical SMILES codes were obtained from the Traditional Chinese Medicine Systems Pharmacology Database and Analysis Platform (TCMSP, https://old.tcmsp-e.com/tcmsp.php), PubChem Database (https://pubchem.ncbi.nlm.nih.gov/) and relevant literatures. Target prediction was performed for each component using the PharmMapper (https://www.lilab-ecust.cn/pharmmapper/index.html), and the PNS target set was established after summarization and deduplication.

#### 2.1.2 Acquisition of Disease-Related Targets

With the keywords of “Alcohol-induced Osteonecrosis of the Femoral Head” and “AIONFH”, the AIONFH-related target genes were retrieved from the GeneCards Database (https://www.genecards.org/), Online Mendelian Inheritance in Man (OMIM, https://www.omim.org/), DisGeNET Database (https://www.disgenet.org/) and Therapeutic Target Database (TTD, http://db.idrblab.net/ttd/). All results were merged and deduplicated to construct the disease target set.

#### 2.1.3 Screening of Drug-Disease Common Targets

The Venn diagram of drug-disease targets was generated using the Jvenn online tool (http://www.bioinformatics.com.cn/static/others/jvenn/example.html) to obtain the intersection targets, which were the potential targets of PNS for the treatment of AIONFH and used for subsequent analysis. The drug and disease targets were imported into Cytoscape 3.9.1 software for visualization to obtain the drug-active component-target network diagram.

#### 2.1.4 Construction and Analysis of Protein-Protein Interaction (PPI) Network

The intersection targets were imported into the STRING database (https://string-db.org/), with the species set as Homo sapiens and the interaction confidence score ≥ 0.4, to obtain PPI network data. The TSV format file was imported into Cytoscape 3.9.1 software for further visualization of its network topological structure.

#### 2.1.5 GO Functional Enrichment and KEGG Pathway Enrichment Analysis

R language software was used to perform GO enrichment analysis and KEGG pathway analysis on the intersection targets. GO analysis included three categories: Biological Process (BP), Cellular Component (CC) and Molecular Function (MF). The significance threshold was set as P < 0.05, and the GO bar plot and KEGG bubble plot were drawn respectively.

#### 2.1.6 Screening of Core Targets by Machine Learning Based on GEO Database

Using the expression matrix of the Gene Expression Omnibus (GEO) database, GSE74089 was selected as the analytical matrix to identify core targets among the intersecting targets via three distinct machine learning algorithms. Logistic regression was implemented with the glmnet package in R, where three-fold cross-validation was employed for feature screening and the importance of variables was assessed on the basis of coefficient magnitudes. Support Vector Machine-Recursive Feature Elimination (SVM-RFE) integrated Support Vector Machine (SVM) with Recursive Feature Elimination (RFE) to optimize variable combinations for target screening. Random Forest models were constructed using the randomForest package in R, with feature importance rankings generated through cross-validation for subsequent target prioritization. Targets identified conjointly by these three algorithms were defined as the final core therapeutic targets for subsequent analysis.

### 2.2 Molecular Docking

The 2D structures of drug small molecule ligands were obtained from the PubChem Database (http://pubchem.ncbi.nlm.nih.gov/), and input into Chem Office software to generate their 3D structures, which were saved as mol2 files. Then the RCSB Protein Data Bank (RCSB PDB, http://www.rcsb.org/) was used to screen the core protein targets. The crystal structure with high resolution was selected as the molecular docking receptor, and Pymol 2.6 software was used to perform dehydration and dephosphorylation on the protein, which was saved as a PDB file. AutoDock 1.5.6 was used to process the structures of proteins and small molecules, including hydrogenation and dehydration of proteins, as well as hydrogenation and torsion angle determination of small molecule ligands, followed by the determination of docking box coordinates. AutoDock Vina software was used for molecular docking to explore the protein-ligand interactions. By comparing the scores of docking results, the optimal conformation of molecular simulation was finally obtained. Discovery Studio 2019 and Pymol 2.6 software were used to visualize the molecular docking results and draw the 2D and 3D analysis diagrams of the interactions between compounds and key residues.

### 2.3 Molecular Dynamics

GROMACS 2023.2 software was used for molecular dynamics simulation in this study. The parameterization of ligands was completed by the GAFF force field in AmberTools22, and the RESP charges were calculated by Gaussian 16w after hydrogenation. The Amber99sb-ildn force field was used to describe the protein in the system, and the simulation was carried out at 300 K and 1 bar with the TIP3P water model, and sodium ions were added to neutralize the system charge **[20]**. The system was first subjected to energy minimization by the steepest descent method with a maximum step size of 50000. Subsequently, the isothermal-isochoric (NVT) ensemble equilibrium and isothermal-isobaric (NPT) ensemble equilibrium were performed in sequence, with a step size of 1 fs for each stage and a total step size of 100,000, corresponding to a simulation time of 100 ps; the V-rescale algorithm was used for temperature coupling, and the Parrinello-Rahman method was used for pressure coupling in the NPT stage. After that, a 50000000-step production run was carried out with a step size of 2 fs, with a total duration of 100 ns. After the simulation was completed, the GROMACS built-in analysis tools were used to process the trajectory, and the Root Mean Square Deviation (RMSD), Root Mean Square Fluctuation (RMSF), Radius of Gyration (Rg) and Solvent Accessible Surface Area (SASA) were calculated. In addition, the binding free energy was calculated by the gmx_MMPBSA tool based on the Molecular Mechanics/Generalized Born Surface Area (MM/GBSA) method, and the energy decomposition analysis was performed using the Generalized Born (GB) solvation model.

### 2.4 Animal Experiment Verification

#### 2.4.1 Animal Preparation

Forty-eight SPF-grade male Sprague-Dawley (SD) rats with a body weight of 200±20 g were purchased from Changsha Tianqin Biotechnology Co., Ltd., with the Experimental Animal Production License No.: SCXK (Xiang) 2019-0014 and Experimental Animal Use License No.: SYXK (Gui) 2022-0001. The rats were raised in a breeding room with a temperature of 20∼25°C, relative humidity of 50%∼65% and a fixed 12 h light/dark cycle per day, and fed with maintenance feed. Humane care was provided to the rats in accordance with the 3R principles (Replacement, Reduction, Refinement) for experimental animal use. The Animal Experimental Ethics Committee Approval No. of Guangxi Zhuang Autonomous Region Academy of Chinese Medical Sciences was 2024093001.

Clinical trial number: not applicable.

#### 2.4.2 Experimental Operations

##### 2.4.2.1 Experimental Grouping and Model Establishment

After 7 days of adaptive feeding, forty-eight SPF-grade male SD rats were randomly divided into the normal control group (8 rats), AIONFH model group (8 rats), compound ossotide positive control group (8 rats), PNS low-dose group (8 rats), PNS medium-dose group (8 rats) and PNS high-dose group (8 rats) using a random number table. Except for the normal control group, the rats in the other groups were given intragastric administration of 46% liquor at a dose of 10 mL/kg once a day for 45 consecutive days to establish the model. After modeling, 2 rats were randomly selected from the AIONFH model group for model identification, and sacrificed by excessive pentobarbital sodium(China National Pharmaceutical Group Shanghai Chemical Reagent Co., Ltd., Batch No.: F20020915) anesthesia. After death, the left femoral head was quickly removed on the animal operating table and fixed in 4% formaldehyde for 48 h, and then Micro CT was used to confirm the successful establishment of the rat AIONFH model.

##### 2.4.2.2 Drug Intervention and Sample Collection

After confirmation of the successful establishment of the AIONFH rat model, the model was maintained and drug administration was initiated. Rats in the normal control group and AIONFH model group were intraperitoneally administered normal saline at a dose of 0.2 mL/kg/day; rats in the Compound Ossotide positive control group were intraperitoneally administered Compound Ossotide Injection (Hebei Zhitong Biopharmaceutical Co., Ltd., National Medicine Approval No. H20052314, Batch No. 0240402) at a concentration of 27.0 mg/mL and a dose of 0.2 mL/kg/day; rats in the low-dose PNS group were intraperitoneally administered low-dose PNS Injection (Beijing Puxitang Biotechnology Co., Ltd., Batch No. C24PA120A) at a concentration of 67.5 mg/mL and a dose of 0.2 mL/kg/day; rats in the medium-dose PNS group were intraperitoneally administered medium-dose PNS Injection at a concentration of 135.0 mg/mL and a dose of 0.2 mL/kg/day; rats in the high-dose PNS group were intraperitoneally administered high-dose PNS Injection at a concentration of 270.0 mg/mL and a dose of 0.2 mL/kg/day. All the above drugs were administered continuously for 45 days. Subsequently, the rats were fasted for 12 hours with free access to water. One hour after the last administration, the rats were anesthetized by intraperitoneal injection of pentobarbital sodium at a dose of 40 mg/kg, blood was collected from the abdominal aorta, and bilateral femoral heads were dissected.

#### 2.4.3 Pathological Staining Analysis of Femoral Head

The left femurs containing the femoral heads of rats in each group were decalcified by the chelating agent (EDTA) method, with the decalcifying solution replaced every 3 days. The decalcification endpoint was defined as the ability of an acupuncture needle to penetrate the bone tissue. Routine pathological paraffin embedding was performed and 4 µm sections were prepared. Tartrate-Resistant Acid Phosphatase (TRAP) staining (Shanghai Gefan Biotechnology Co., Ltd., Cat. No.: M071, Batch No.: 20250317), Hematoxylin-Eosin (HE) staining (Beijing Leagene Biotechnology Co., Ltd., Batch No.: 0509A22), Alizarin Red staining (Shanghai Gefan Biotechnology Co., Ltd., Cat. No.: M040, Batch No.: 20250317) and Alkaline Phosphatase (AKP) staining (Shanghai Gefan Biotechnology Co., Ltd., Cat. No.: M039, Batch No.: 20250317) were conducted to observe the pathological changes of the femoral heads.

#### 2.4.4 Detection of Target Protein Expression Localization and Level by Immunofluorescence Staining

The paraffin sections of the right femoral head tissue of rats in each group were dewaxed to water and stained by the routine immunofluorescence method. The distribution and expression of HSD11B1 (1:500) protein in the femoral head were observed under a microscope, and images were collected under a fluorescence microscope.

#### 2.4.5 Detection of Target Gene Expression by qPCR

Total RNA was extracted from the right femoral head tissue of rats by the Trizol method and reverse-transcribed into cDNA. Then quantitative real-time polymerase chain reaction (qPCR) was used to quantitatively detect the mRNA expression of Runx2, CTSK and HSD11B1 in the femoral head. The primer sequences and experimental conditions for quantitative PCR are detailed in Table 1.

#### 2.4.6 Statistical Methods

All experimental measurement data were expressed as mean ± standard deviation (±s) and analyzed using SPSS 16.0 software. Normality test verified that the data conformed to a normal distribution. One-way analysis of variance (ANOVA) was used for comparisons among multiple groups. Bonferroni correction was adopted for pairwise comparisons when variances were homogeneous, while Tamhane’s T2 test was used when variances were heterogeneous. A P value < 0.05 indicated statistical significance. In statistical annotation, P < 0.05 was denoted as “*”, P < 0.01 as “**”, P < 0.001 as “***”, P < 0.0001 as “****”, and P ≥ 0.05 as “ns”.

## 3 Results

### 3.1 Network Pharmacology

#### 3.1.1 PNS Active Components and Corresponding Targets

Based on the TCMSP and SwissTargetPrediction databases and combined with relevant literature research, the potential targets corresponding to the effective components in PNS were screened out. A total of 5 main active components were identified, including Notoginsenoside R1 (NG-R1), Ginsenoside Rb1 (G-Rb1), Ginsenoside Rg1 (G-Rg1), Ginsenoside Re (G-Re) and Ginsenoside Rd (G-Rd). A total of 127 potential targets of PNS were finally obtained after data integration and deduplication **[21-24]**.

#### 3.1.2 AIONFH Target Set and Its Intersection Targets with PNS

AIONFH-associated targets were obtained from the GeneCards, OMIM, DisGeNET and TTD databases. All records were merged and deduplicated to establish a disease target set comprising 672 unique targets. As shown in Figure 1A, the intersection between the disease target set and the potential target set of PNS was identified and visualized via the Jvenn platform to generate a drug-disease target Venn diagram, yielding a total of 18 overlapping targets. These drug and disease targets were imported into Cytoscape 3.9.1 for visualization, and the PNS-AIONFH drug-active component-target network diagram was constructed as illustrated in Figure 1B.

**Figure 1.**
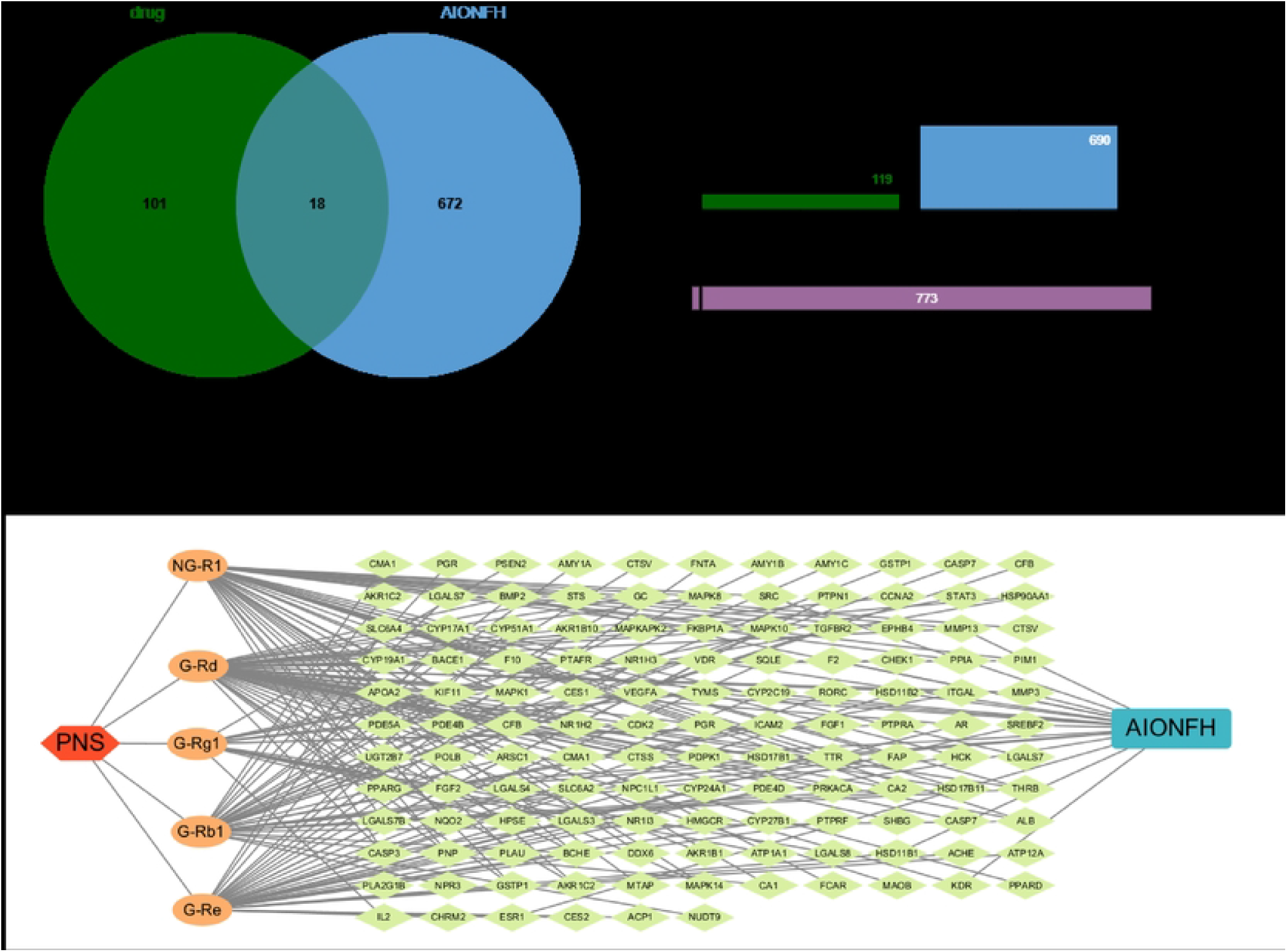
Venn diagram of drug-disease targets and network diagram of drug-active ingredient-action targets for PNS and AIONFH. Panel A is the Venn diagram of drug-disease targets for Panax Notoginseng Saponins (PNS)-related genes and Alcohol-induced Osteonecrosis of the Femoral Head (AIONFH)-related genes, which clearly shows the 18 overlapping targets obtained by intersecting the potential targets of PNS and the disease-related targets of AIONFH. Panel B is the visualized network diagram of drug-active ingredient-action targets between PNS and AIONFH constructed by Cytoscape 3.9.1 software, which displays the interaction relationships among PNS, its main active ingredients and the overlapping targets of the disease.

#### 3.1.3 Construction of PPI Network for 18 Intersection Targets

As shown in Figure 2A, the 18 intersection targets were uploaded to the STRING database for the construction of a PPI network, which comprised 18 nodes and 55 edges with an average node degree of 6.11. F2 was excluded from subsequent analyses owing to its isolation from the primary network.The resulting network contained 17 nodes and 55 edges, and the PPI network was further optimized using Cytoscape 3.9.1, as illustrated in Figure 2B.

**Figure 2.**
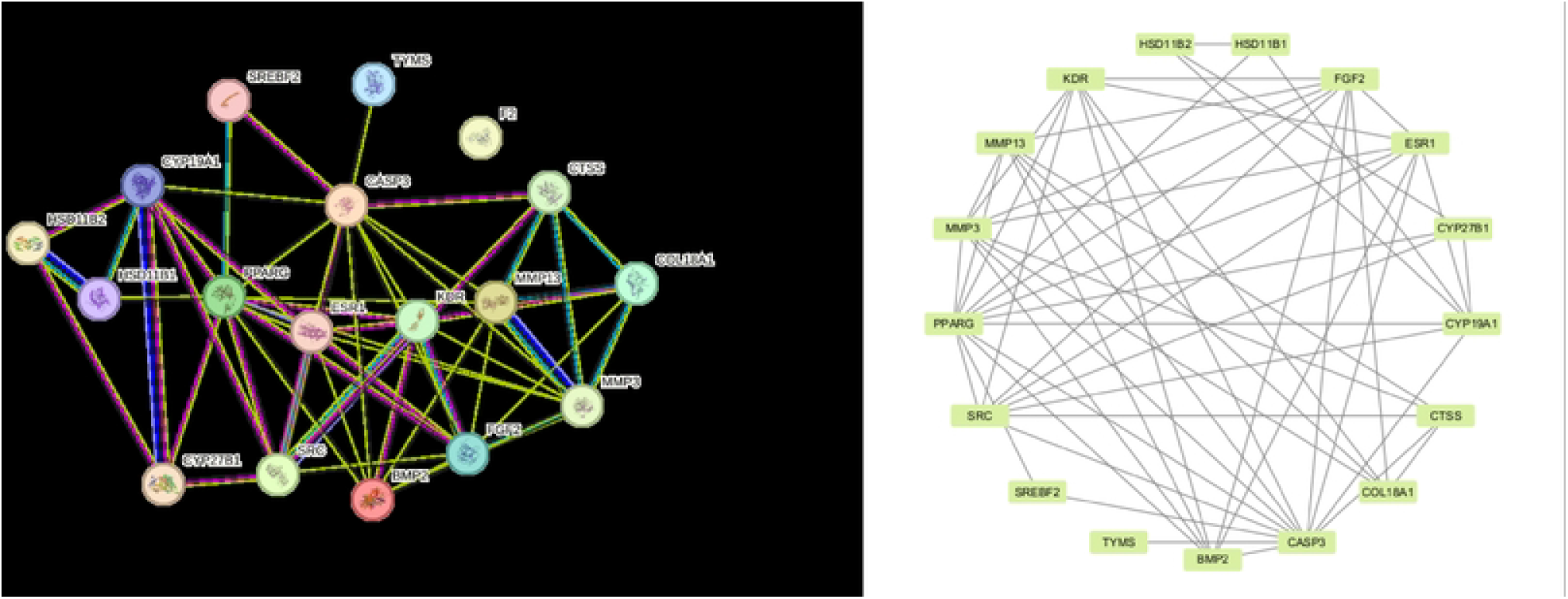
Protein-protein interaction (PPI) network diagrams of the overlapping targets. Panel A is the PPI network diagram of the 18 overlapping targets constructed by the STRING database, which contains 18 nodes and 55 edges with an average node degree of 6.11. Panel B is the optimized PPI network diagram processed by Cytoscape 3.9.1 software, in which the F2 target separated from the main network is excluded, and the remaining network consists of 17 nodes and 55 edges for subsequent in-depth analysis.

#### 3.1.4 GO Enrichment Analysis and KEGG Pathway Enrichment Analysis

R was employed to conduct Gene Ontology (GO) functional enrichment analysis and Kyoto Encyclopedia of Genes and Genomes (KEGG) pathway enrichment analysis on the 18 intersection targets. Using a significance threshold of P < 0.05, GO enrichment analysis revealed 1042 significant biological process (BP) terms, 14 significant cellular component (CC) terms, and 109 significant molecular function (MF) terms, from which the GO bar plot was generated as shown in Figure 3A. KEGG pathway enrichment analysis yielded 38 significant pathways in total; the top 10 pathways, ranked by ascending P values, were selected for the generation of the KEGG bubble plot as illustrated in Figure 3B.

**Figure 3.**
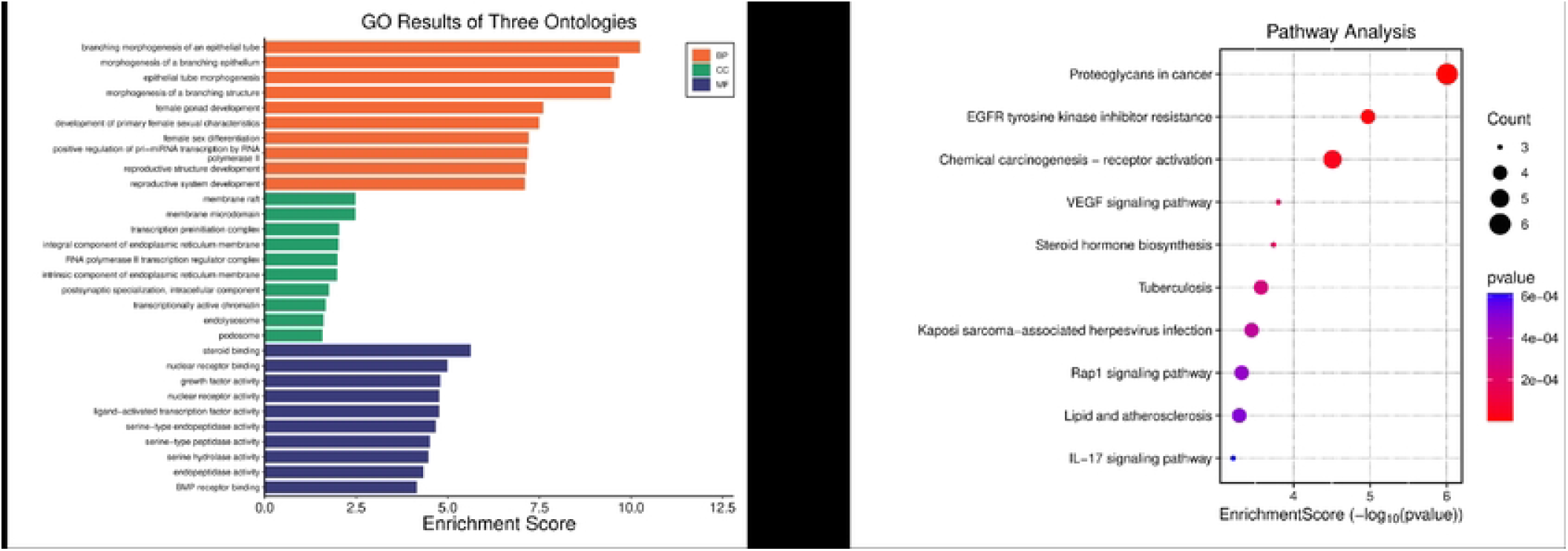
Histogram of Gene Ontology (GO) enrichment analysis and bubble plot of Kyoto Encyclopedia of Genes and Genomes (KEGG) pathway enrichment analysis. Based on the 18 overlapping targets with the significance threshold of p<0.05, Panel A is the GO enrichment analysis histogram, which includes 1042 significant entries for biological processes, 14 for cellular components and 109 for molecular functions. Panel B is the KEGG pathway enrichment analysis bubble plot, in which the top 10 significant pathways are selected for plotting according to the ascending order of p values from the total 38 significant pathways obtained by the analysis.

#### 3.1.5 Screening of Core Targets by Machine Learning Based on GEO Database

To further screen the potential core therapeutic targets of PNS for AIONFH, dataset GSE74089 from the Gene Expression Omnibus database was utilized as the analytical matrix. Three machine learning algorithms, including logistic regression, SVM-RFE and random forest, were adopted to screen the 18 intersection targets. As shown in Figure 4A, targets consistently identified by all three approaches were designated as the final core therapeutic targets and visualized in a Venn diagram. Six core targets were ultimately identified, including FGF2, HSD11B1, BMP2, MMP13, TYMS and VEGFA. Figure 4B depicts the box scatter plot for inter-group gene expression comparison, which delineates the expression profiles of the six core target genes in dataset GSE74089. Comparison of expression levels between the case group and the control group uncovered significant differences between the two groups, thereby validating the potential value of these targets as key molecules. Figure 4C illustrates the importance distribution of SVM-RFE, random forest and logistic regression algorithms in the screening model. Figure 4D and Figure 4E characterize cross-algorithm feature importance and ranking attenuation via line charts and heat maps, respectively.

**Figure 4.**
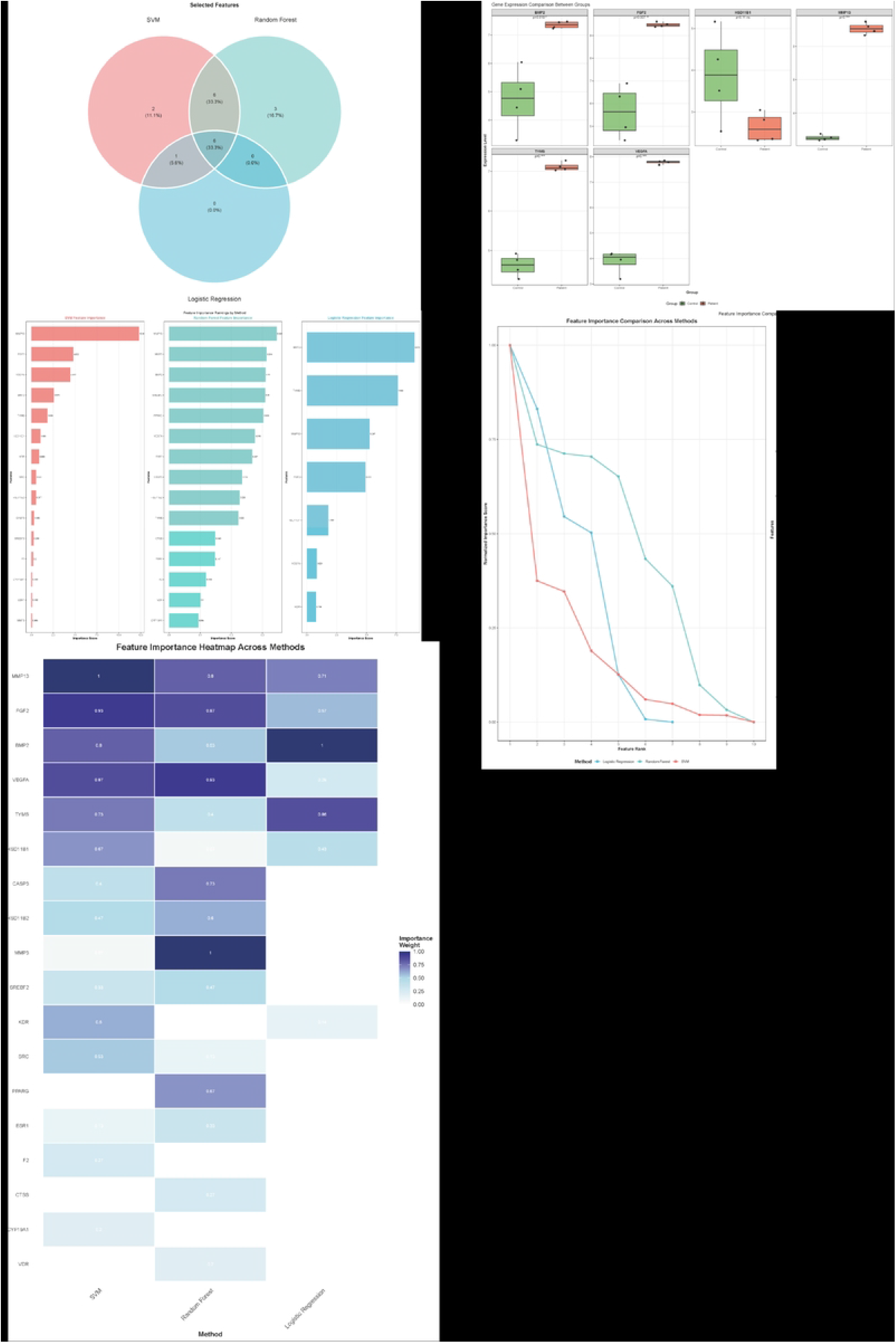
Identification of core targets by machine learning algorithms. Panel A is the Venn diagram of core genes, which screens out the common targets of the three machine learning algorithms as the final core therapeutic targets for PNS in the treatment of AIONFH. Panel B is the box scatter plot of inter-group comparison of core gene expression, which shows the expression profiles of the core target genes in the GSE74089 dataset and the significant expression differences between the case group and the control group. Panel C is the histogram of the importance distribution of the three machine learning algorithms (Logistic regression, SVM-RFE, Random Forest) in the screening model. Panel D is the line chart of cross-algorithm comparison of feature importance, and Panel E is the heat map of cross-algorithm comparison of feature importance and ranking attenuation, which jointly reflect the importance of each target in different screening algorithms.

### 3.2 Molecular Docking

Molecular docking studies were conducted to further explore the interaction between PNS and AIONFH-related core proteins, aiming to elucidate the potential mechanism of PNS in the treatment of AIONFH. Machine learning analysis identified 6 core targets: FGF2, HSD11B1, BMP2, MMP13, TYMS and VEGFA. Then 5 active components of PNS (NG-R1, G-Rb1, G-Rg1, G-Re and G-Rd) were subjected to molecular docking analysis with these 6 targets. The molecular docking results of PNS and target proteins are shown in Table 2, the heat map is shown in Figure 5, and the 6 pairs with the best molecular docking binding energy are shown in Figure 6 after visualization. The results showed that the docking scores of all PNS-target protein complexes were lower than -5.0 kcal/mol. According to the established criteria, a lower docking score indicates a stronger binding affinity; a score less than -5.0 kcal/mol indicates potential binding activity, and a score less than -7.0 kcal/mol indicates a strong binding affinity **[25]**. Notably, HSD11B1 exhibited the strongest binding affinity with G-Re, an active component of PNS, with a score of -12.4 kcal/mol.

**Figure 5.**
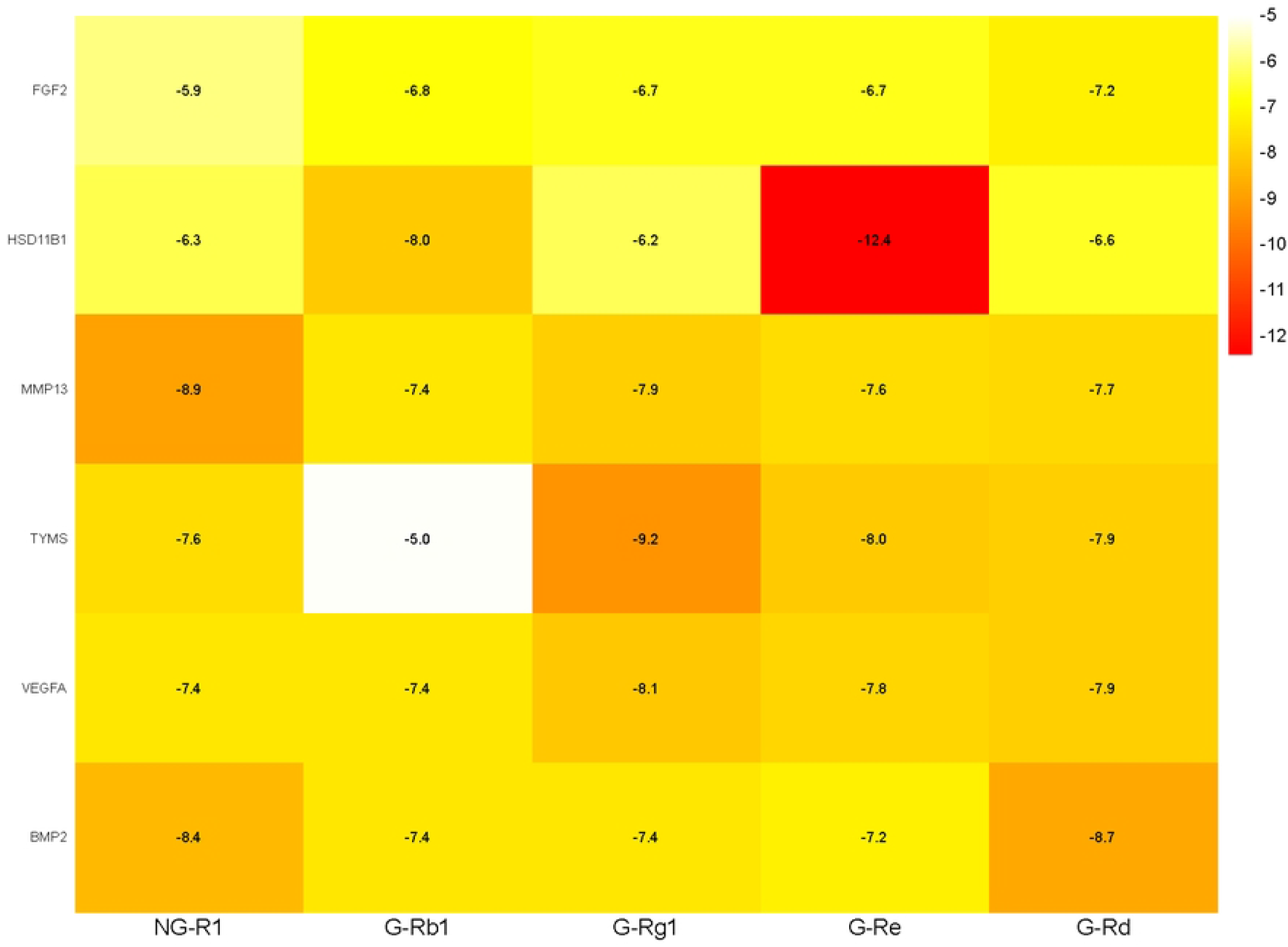
Heat map of molecular binding energy between PNS active ingredients and core target proteins. This heat map shows the molecular docking binding energy values (in kcal/mol) of five main active ingredients of PNS with six core target proteins of AIONFH. The color gradient of the heat map indicates the magnitude of the absolute value of the binding energy, with redder color representing a larger absolute value of binding energy, and a lower binding energy value indicating a stronger binding affinity between the small molecule and the protein.

**Figure 6.**
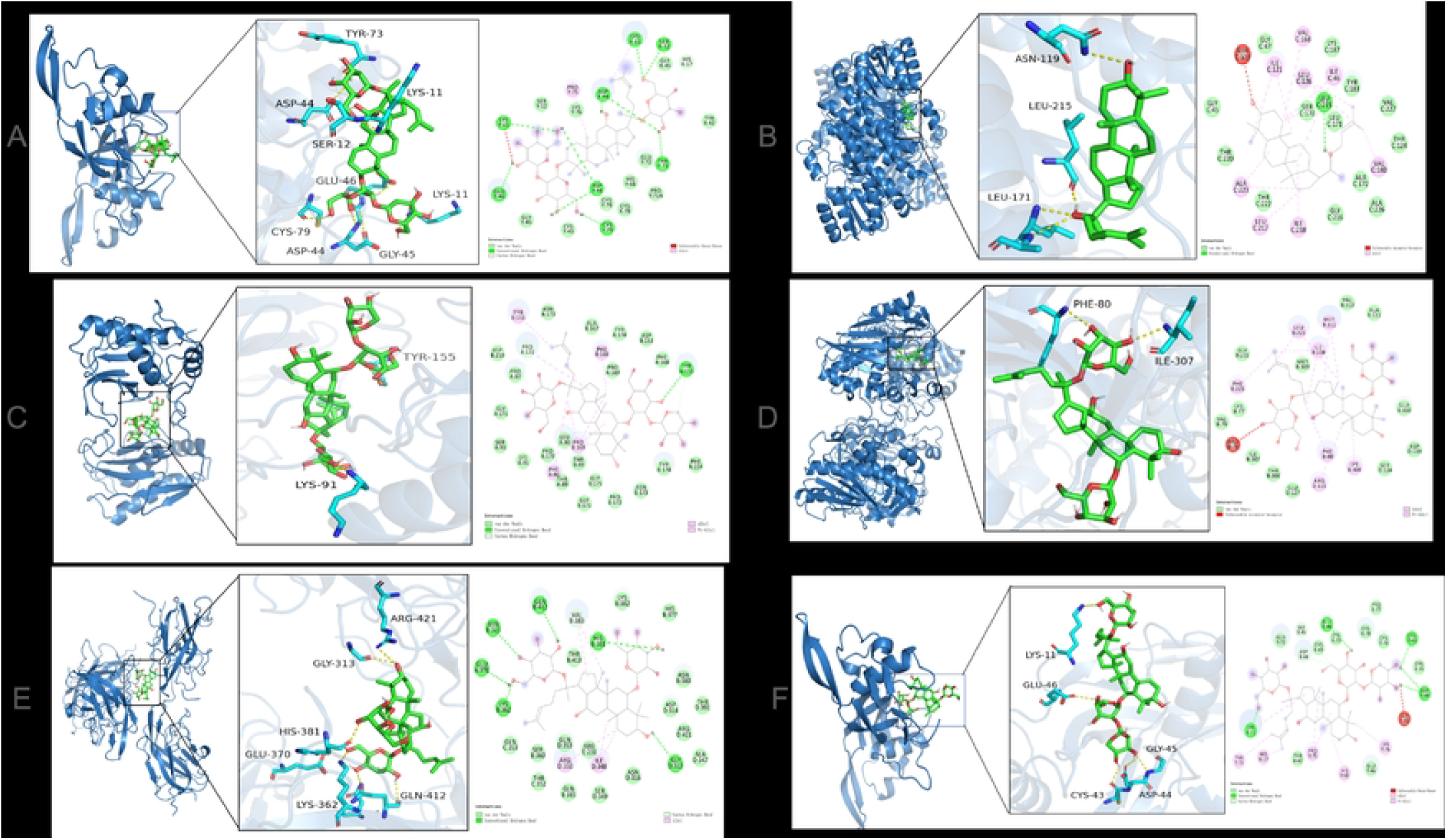
Visualization of molecular docking results between core target proteins and PNS active ingredients. This figure displays the two-dimensional visualization results of the six pairs of protein-ligand complexes with the optimal molecular docking binding energy. Panel A shows the docking result of BMP2 and Ginsenoside Rd (G-Rd); Panel B shows the docking result of HSD11B1 and Ginsenoside Re (G-Re); Panel C shows the docking result of MMP13 and Notoginsenoside R1 (NG-R1); Panel D shows the docking result of TYMS and Ginsenoside Rg1 (G-Rg1); Panel E shows the docking result of VEGFA and G-Rg1; Panel F shows the docking result of BMP2 and NG-R1, all of which clearly present the key amino acid residues involved in the binding interaction.

### 3.3 Molecular Dynamics

Among the molecular docking results of all target proteins and PNS active components, 11β-Hydroxysteroid Dehydrogenase Type 1 (HSD11B1) and G-Re showed the strongest binding energy. Molecular dynamics analysis was performed on the molecular docking results of HSD11B1 and G-Re, and the results are shown in Figure 7.

**Figure 7.**
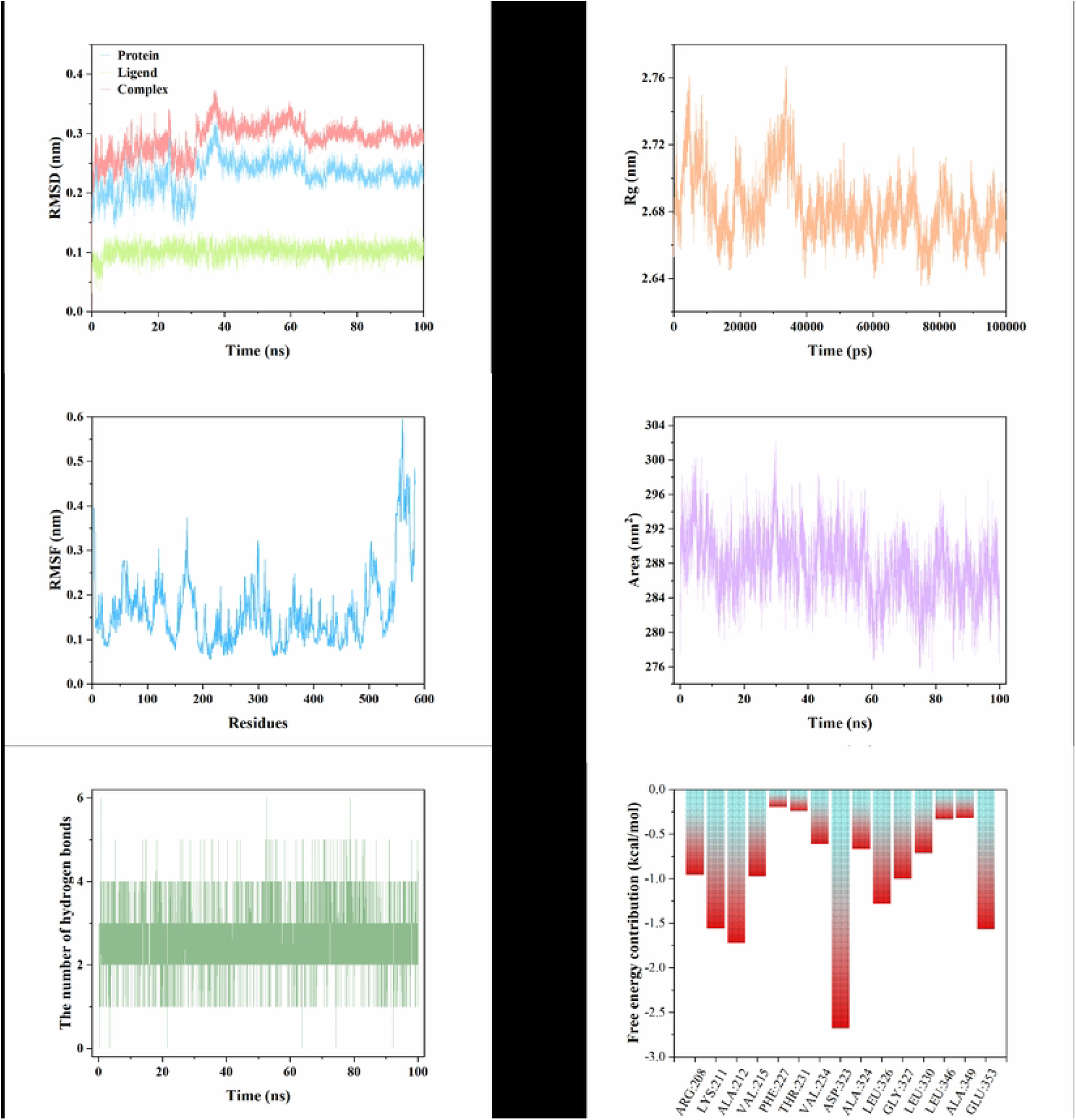
Molecular dynamics simulation results of the HSD11B1-G-Re complex within 100 ns. This figure evaluates the conformational stability of the HSD11B1-G-Re complex through multiple key indicators of molecular dynamics simulation. Panel A is the root mean square deviation (RMSD) plot reflecting the conformational offset degree of the complex relative to the initial state; Panel B is the radius of gyration (Rg) plot characterizing the structural compactness of the protein; Panel C is the root mean square fluctuation (RMSF) plot representing the flexibility of each amino acid residue of the protein; Panel D is the solvent accessible surface area (SASA) plot quantifying the surface area of the molecule exposed to the solvent; Panel E is the plot of the number of hydrogen bonds between HSD11B1 and G-Re during the simulation; Panel F is the plot of residue free energy contribution, which identifies the key amino acid residues that play a core role in the stable binding of the complex.

RMSD is a core index for evaluating the conformational stability of protein-ligand complexes, reflecting the deviation degree of atoms relative to the initial conformation. Generally, an RMSD value maintained at 0.1∼0.5 nm indicates a stable system [26]. As shown in Figure 5A, the RMSD value of the G-Re-HSD11B1 complex was consistently lower than 0.4 Å (0.04 nm) during the entire simulation period (0∼100 ns) with an extremely small fluctuation range (< 0.05 Å). The RMSD curves of the receptor protein and the protein complex were highly coincident without an obvious upward trend, indicating that no significant conformational rearrangement occurred after the binding of G-Re to HSD11B1, the overall structure of the complex had high thermodynamic stability, and no drift phenomenon of the ligand occurred in the protein binding pocket.

Rg is used to describe the structural compactness of proteins in three-dimensional space, and the stability of its value directly reflects the folding state of the protein backbone [26]. As shown in Figure 7B, during the 100 ns simulation period, the Rg value of the complex was consistently maintained at 2.64∼2.76 Å with a fluctuation range of less than 0.05 Å and no significant increase or decrease trend. This result indicated that the binding of G-Re to HSD11B1 did not cause loosening or excessive folding of the protein structure, and the protein backbone always maintained a stable compact conformation, further verifying the overall structural stability of the complex.

RMSF is used to characterize the flexibility of each amino acid residue of the protein; a low RMSF value corresponds to a structurally stable core region, while a high value indicates a highly flexible loop region or terminal [26]. As can be seen from Figure 7C, in the G-Re-HSD11B1 complex, the RMSF values of most residues were lower than 0.4 Å, only a few terminal residues had a slightly higher fluctuation range, and the maximum value did not exceed 0.6 Å. This indicated that ligand binding did not damage the rigidity of the protein core structure, and the stable conformation of the binding pocket region provided a structural basis for the continuous interaction between the two.

SASA is used to quantify the area of the molecular surface exposed to the solvent, and its changes can reflect the effect of ligand binding on the protein conformation and the microenvironment of the binding pocket [26]. As can be seen from Figure 7D the SASA value of the complex was maintained in the range of 276∼304 arbitrary units during the simulation with a fluctuation range of less than 5% and no significant increase or decrease trend. This indicated that after G-Re binding, the exposure degree of hydrophobic groups on the HSD11B1 surface did not change significantly, and the microenvironment of the binding pocket was in a stable state without hydrophobic imbalance on the protein surface caused by ligand binding, which guaranteed the thermodynamic stability of the complex.

Hydrogen bonds are key non-covalent interactions maintaining the stability of protein complexes, and their quantity and persistence directly affect the binding affinity **[26]**. As shown in Figure 7E, the number of hydrogen bonds between G-Re and HSD11B1 remained stable during the entire simulation period, mostly maintained at 2∼3, without a continuous decrease or sharp drop. The stable hydrogen bond interaction indicated that G-Re could form continuous hydrogen bonds with specific residues (such as ALA172 and ILE46) in the HSD11B1 binding pocket, which provided important support for the stable binding of the complex and was consistent with the molecular docking results.

Residue free energy decomposition based on the MM/GBSA method can clarify the contribution of a single amino acid residue to ligand binding, and a significant negative value indicates that the residue is a key functional site **[27-28]**. As can be seen from Figure 7F and Table 3, in the G-Re-HSD11B1 protein complex, the free energy contribution values of 15 key residues were all negative, among which ASP:323 had the lowest contribution energy (-2.679) and was the core residue for the stable binding of the ligand; ALA:212 (-1.72), GLU:353 (-1.568) and LYS:211 (-1.559) had the next lowest contribution values and were strong binding residues; the absolute values of the contribution energy of residues such as PHE:227 (-0.197) and THR:231 (-0.243) were small, with limited effects on the binding stability. These results indicated that the binding of G-Re to HSD11B1 mainly depended on the synergistic effect of key residues such as ASP:323 and ALA:212, involving hydrogen bonds, electrostatic interactions and hydrophobic interactions, further revealing the molecular mechanism of the specific binding between the two.

In summary, the molecular dynamics simulation results confirmed that the complex formed by G-Re and HSD11B1 had good conformational stability and binding specificity. The complex exhibited low fluctuation characteristics in core indexes such as RMSD, Rg and SASA, and the stable hydrogen bond interaction and strong free energy contribution of key residues jointly guaranteed the continuous specific binding between the ligand and the protein. This provided reliable microdynamic evidence for PNS to target and inhibit HSD11B1 activity and improve the bone marrow microenvironment of AIONFH through G-Re.

### 3.4 Animal Experiment Verification

#### 3.4.1 Results of Femoral Head Pathological Staining

In this study, TRAP, HE, Alizarin Red and ALP staining were used to analyze the therapeutic effect of PNS on AIONFH from multiple links of bone metabolism, as detailed in Figure 8.

**Figure 8.**
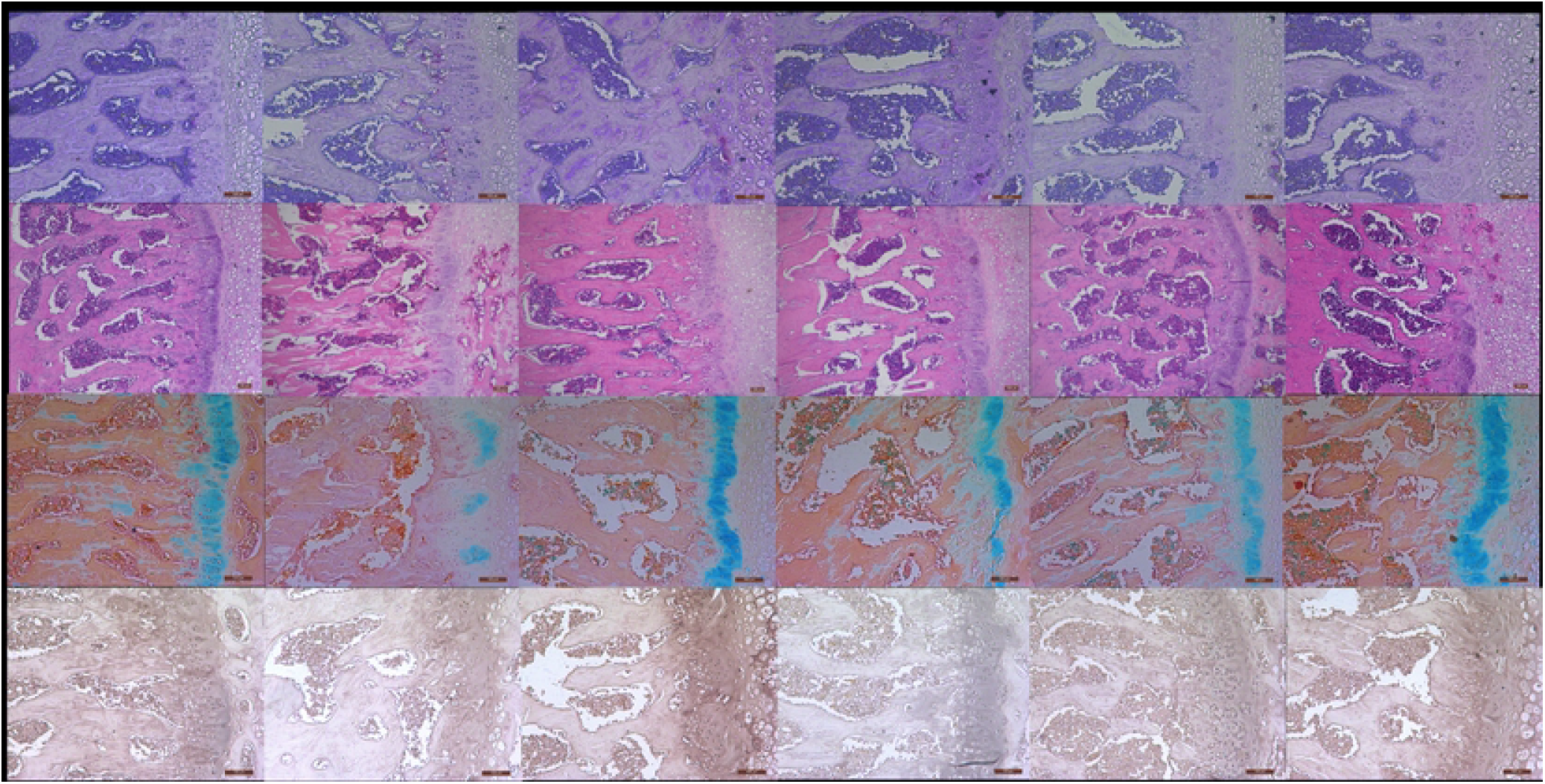
Pathological staining of femoral heads in each experimental group of rats. This figure shows the results of tartrate-resistant acid phosphatase (TRAP) staining, hematoxylin-eosin (HE) staining, Alizarin Red staining and alkaline phosphatase (AKP) staining of rat femoral heads in different groups, with the magnification of ×100 for TRAP, Alizarin Red and AKP staining, and ×50 for HE staining. The groups are labeled as Normal Control (NC), Alcohol-induced Osteonecrosis of the Femoral Head model (AIONFH), Compound Ossotide Positive Control (PC), Low-dose PNS, Medium-dose PNS and High-dose PNS, which intuitively reflect the therapeutic effect of PNS on the pathological structure of femoral heads in AIONFH rats with a dose-dependent manner.

Under TRAP staining (×100), the distribution and activity of osteoclasts in the normal control group maintained a physiological steady state **[29]**. Osteoclasts were overactivated with abnormally enhanced bone resorption in the AIONFH model group, while the overactive state of osteoclasts in each PNS dose group was gradually inhibited with the increase of dose, and the high-dose group was close to the performance of the normal control group, suggesting that PNS could effectively regulate the balance of bone resorption.

Under HE staining (×100), the bone tissue structure in the normal control group was intact, and the morphology of osteocytes and trabecular bone as well as the distribution of marrow adipocytes were orderly **[30]**. The trabecular bone in the model group was sparse and fractured, marrow adipocytes proliferated abnormally, osteocytes were arranged disorderly, the rate of empty lacunae increased, and bone tissue was damaged significantly; after PNS intervention, the morphology of trabecular bone was gradually repaired, the abnormal proliferation of marrow adipocytes was limited, the bone tissue structure of the medium- and high-dose groups was close to normal, and osteocytes were arranged regularly, indicating that PNS could promote the repair of bone tissue structure and optimize the bone marrow microenvironment.

In Alizarin Red staining (×100), calcium deposition in the normal control group was uniform with clear staining **[31]**. Calcium deposition in the model group was sharply reduced with light staining, and the bone mineralization process was hindered; after PNS intervention, calcium deposition was gradually improved with the increase of dose, and the range and intensity of calcium deposition in the high-dose group were close to normal, indicating that PNS could effectively promote bone tissue mineralization and restore the balance of calcium metabolism.

The results of ALP staining (×100) showed that the activity of osteoblasts in the normal control group was normal with significant positive staining **[32]**. The activity of osteoblasts in the model group was inhibited with weakened positive staining; the activity of osteoblasts in each PNS dose group was gradually enhanced with the increase of dose, and the high-dose group had obvious positive staining with well-recovered osteogenic function, confirming that PNS could activate osteoblasts and enhance bone formation ability.

#### 3.4.2 Immunofluorescence Staining of HSD11B1 Protein

As shown in Figure 9, immunofluorescence staining demonstrated that HSD11B1 protein was predominantly localized in the cytoplasm of femoral head tissue. DAPI staining showed blue cell nuclei, and HSD11B1 protein showed green fluorescence. The fluorescence intensity of HSD11B1 protein in the femoral head tissue of the normal control group (NC) was weak. the fluorescence intensity of HSD11B1 protein in the AIONFH model group was significantly enhanced compared with the NC group; after PNS intervention, the fluorescence intensity of HSD11B1 protein in each dose group gradually decreased with the increase of dose. The results indicated that the expression of HSD11B1 protein in the femoral head tissue of AIONFH model rats was significantly up-regulated, and PNS could dose-dependently inhibit the abnormally high expression of HSD11B1 protein in the AIONFH model.

**Figure 9.**
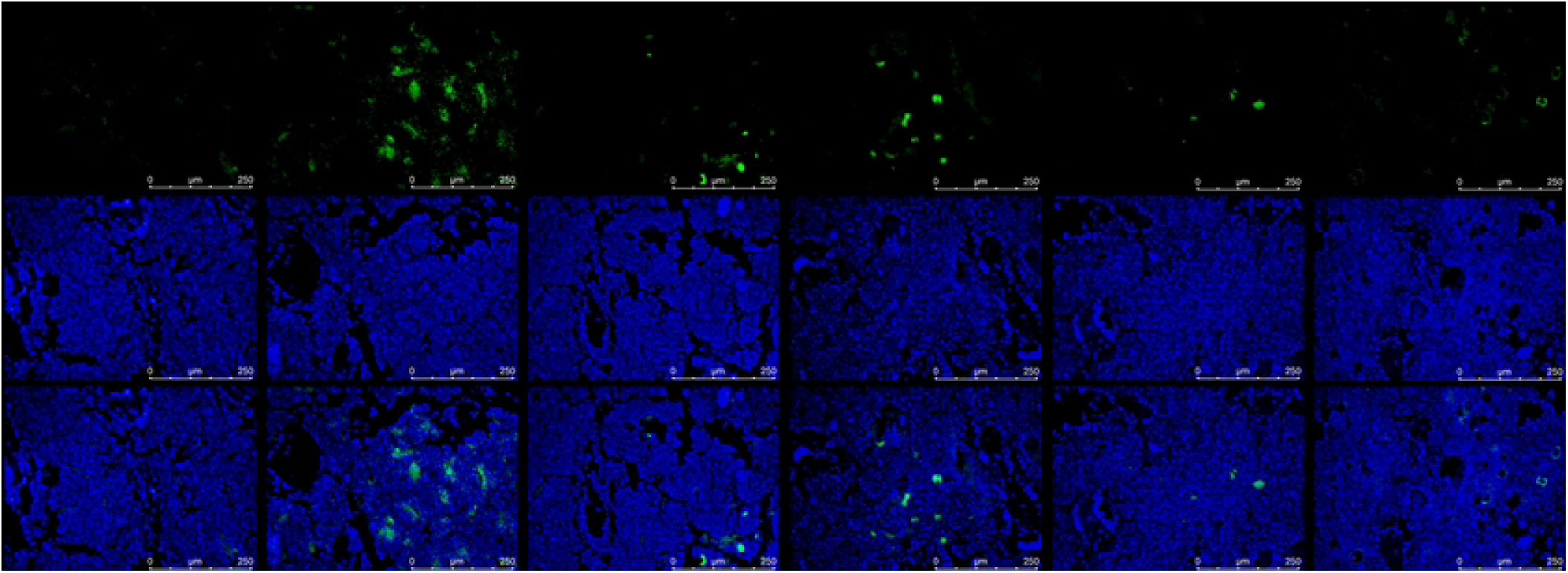
Immunofluorescence staining of HSD11B1 protein in rat femoral head tissues. This figure shows the immunofluorescence staining results of HSD11B1 protein in femoral head tissues of each experimental group (NC, AIONFH, PC, Low-dose PNS, Medium-dose PNS, High-dose PNS). DAPI staining labels the cell nucleus in blue, and HSD11B1 protein is labeled with green fluorescence, which is mainly localized in the cytoplasm. The fluorescence intensity directly reflects the expression level of HSD11B1 protein, showing the dose-dependent inhibitory effect of PNS on the abnormally high expression of HSD11B1 protein in AIONFH model rats.

### 3.5 Results of HSD11B1, Runx2 and CTSK Gene Expression Detected by qPCR

As shown in Figure 10, qPCR analyses revealed that the relative expression level of the HSD11B1 gene was significantly down-regulated in the positive control (PC) group relative to the alcohol-induced osteonecrosis of the femoral head (AIONFH) model group, with an extremely significant difference (P<0.0001).No significant difference in the expression level of HSD11B1 was observed between the PNS low dose group and the AIONFH group (ns), while the relative expression levels of HSD11B1 in the PNS medium and high dose groups were both extremely significantly lower than those in the AIONFH group (P<0.0001).Runt-related transcription factor 2 (Runx2) is a key transcription factor for osteogenic differentiation **[33]**. The relative expression level of Runx2 in the PC group was extremely significantly decreased compared with that in the AIONFH group (P<0.0001). No significant difference in the expression level of Runx2 was found between the PNS low dose group and the AIONFH group (ns), while the relative expression levels of Runx2 in the PNS medium and high dose groups were both extremely significantly lower than those in the AIONFH group (P<0.0001).Cathepsin K (CTSK) is one of the core markers in the study of osteoclast function and a key effector molecule in the process of osteoclast-mediated bone resorption **[34]**. The relative expression level of CTSK in the AIONFH group was significantly higher than that in the NC group. Further intergroup analysis showed that although the expression levels of CTSK in the PC group and the PNS low, medium, and high dose groups were all lower than those in the AIONFH group, there were no statistically significant differences between each group and the AIONFH group, as well as between the groups (ns).

**Figure 10.**
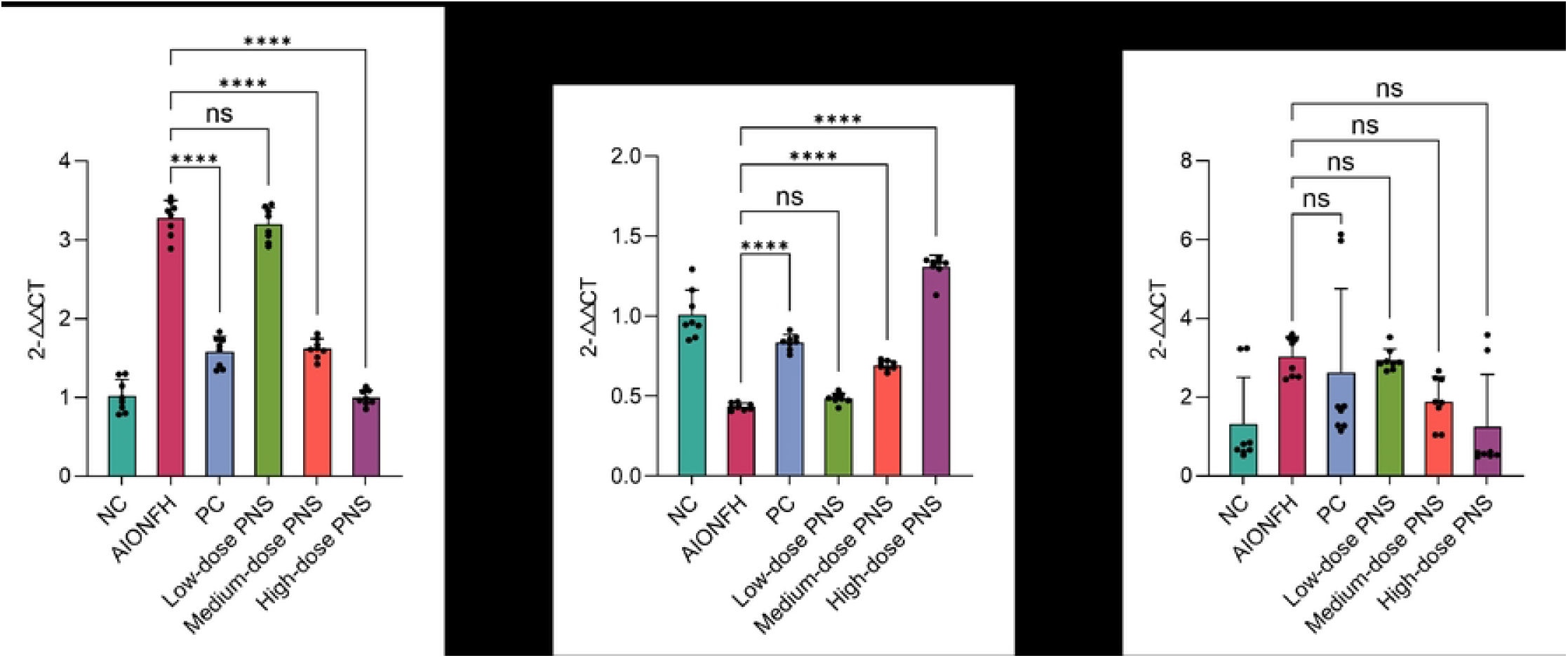
Relative mRNA expression levels of HSD11B1, Runx2 and CTSK detected by quantitative real-time polymerase chain reaction (qPCR). This figure presents the qPCR detection results of the relative mRNA expression levels of HSD11B1, Runt-related transcription factor 2 (Runx2) and Cathepsin K (CTSK) in femoral head tissues of each group (NC, AIONFH, PC, Low-dose PNS, Medium-dose PNS, High-dose PNS). The results show the abnormal expression of the three genes in the AIONFH model group and the dose-dependent reversal effect of PNS on the abnormal gene expression, that is, inhibiting the expression of HSD11B1 and CTSK and promoting the expression of Runx2.In the figure, P<0.001 is marked as ‘****’, and P≥0.05 is marked as ‘ns’.

The above results indicated that the expressions of HSD11B1 and CTSK genes were abnormally up-regulated, and the expression of Runx2 gene was significantly down-regulated in the femoral head tissue of AIONFH model rats. PNS intervention could dose-dependently reverse the abnormal expressions of HSD11B1 and Runx2, among which the inhibitory effect on HSD11B1 and the promoting effect on Runx2 expression were statistically significant at medium and high doses, while the down-regulation of CTSK only showed a trend of change. It is suggested that PNS may regulate the osteoblast-osteoclast balance mainly by inhibiting HSD11B1 expression and promoting Runx2 expression, thereby exerting a therapeutic effect on AIONFH.

## 4 Discussion

In this study, network pharmacology, machine learning, molecular docking, molecular dynamics simulation and animal experiments were integrated to systematically elucidate the core mechanism of PNS in the treatment of AIONFH from the multi-dimensional perspective of “component-target-pathway-pathological phenotype”, thereby providing a scientific basis for its clinical translation.

The pathological essence of AIONFH is the interruption of femoral head blood supply, osteocyte apoptosis and bone matrix destruction caused by long-term alcohol exposure **[35]**. Long-term alcohol intake can elevate the activity of HSD11B1, which promotes the conversion of inactive corticosterone to active cortisol, thereby exacerbating the abnormal proliferation of bone marrow adipocytes and compressing the space of intraosseous capillaries **[36-37]**. Meanwhile, it inhibits the expression of angiogenic factors such as VEGFA and FGF2, forming a vicious cycle of “ischemia-osteonecrosis” **[38-41]**. In addition, alcohol can activate MMP13-mediated bone matrix degradation and inhibit osteoblast differentiation through the positive regulation of BMP2, which further impairs the structural stability of bone tissue **[42-45]**. Network pharmacology screening in this study identified 18 common potential targets between the core active components of PNS and AIONFH. Gene Ontology (GO) functional enrichment analysis revealed that these common potential targets are mainly involved in key biological processes such as osteoblast differentiation, which is highly consistent with the “ischemia-osteonecrosis” pathological process of AIONFH. KEGG pathway analysis further indicated that the VEGF pathway is the core pathway for PNS to exert its therapeutic effects, and this pathway can directly regulate the proliferation and lumen formation of vascular endothelial cells, providing a critical pathway basis for improving the ischemic microenvironment of the femoral head. The above targets were further screened by three machine learning algorithms including Logistic regression, SVM-RFE and random forest, and six core targets were finally identified, namely FGF2, HSD11B1, BMP2, MMP13, TYMS and VEGFA. This screening strategy significantly improved the reliability and clinical relevance of the identified targets.

Molecular docking technology is a core method to verify the binding specificity between drug components and targets. In this study, molecular docking analysis was performed between the five active components of PNS and the six core targets, and the results showed that the binding energy of all component-target complexes was lower than -5.0 kcal/mol, reaching the threshold of potential binding activity. Notably, Ginsenoside Re (G-Re) had the lowest binding energy with HSD11B1 (-12.4 kcal/mol), which was far below the criterion of -7.0 kcal/mol for strong binding affinity, suggesting that this combination constitutes the key molecular basis for PNS to exert its pharmacological effects. To further verify the conformational stability of this complex, 100 ns molecular dynamics simulation was conducted using Gromacs software, and the results demonstrated that the RMSD of the complex was consistently below 0.04 nm with minimal fluctuations throughout the simulation. The Rg was stably maintained in the range of 2.64∼2.76 Å, and the fluctuation of the SASA was less than 5%. Meanwhile, 2 to 3 hydrogen bonds were stably maintained between G-Re and HSD11B1. The above indicators all confirmed that G-Re and HSD11B1 can form a stable binding conformation, and no drift of the ligand was observed in the protein binding pocket. The results of residue free energy decomposition further identified that key residues such as ASP:323 and ALA:212 provide the core support for the stable binding of the complex, which reveals the molecular mechanism of PNS targeting HSD11B1 at the atomic level.

Animal experiments are the gold standard for verifying the in vivo therapeutic effects of drugs. In this study, a rat model of AIONFH was established, and a series of pathological staining techniques were used to analyze the therapeutic effects of PNS from multiple aspects of bone metabolism, which confirmed the dose-dependent therapeutic effect of PNS on AIONFH. HE staining results showed that PNS can repair the damaged trabecular bone structure, reduce the abnormal proliferation of bone marrow adipocytes, decrease the rate of empty osteocyte lacunae and restore the structural integrity of femoral head tissue. TRAP staining confirmed that PNS can effectively inhibit the excessive activation of osteoclasts, reverse the pathological state of hyperactive bone resorption and precisely regulate the balance of bone resorption. AKP staining further verified that PNS can significantly promote calcium salt deposition in bone tissue, enhance osteoblast activity, accelerate the bone mineralization process and reconstruct the physiological balance between bone formation and bone resorption. To further elucidate the underlying mechanism, immunofluorescence staining and qPCR were used to verify the regulatory effect of PNS on the core targets at the protein and transcriptional levels, respectively. In this study, the protein and gene expressions of HSD11B1 in the femoral head tissue of AIONFH model rats were significantly up-regulated, and PNS intervention led to a significant dose-dependent reduction in its expression. Meanwhile, PNS could effectively reverse the down-regulation of Runx2, a key transcription factor for osteogenic differentiation, thus achieving the precise regulation of bone metabolic balance. As a key enzyme regulating the local glucocorticoid metabolism in the femoral head, HSD11B1 can catalyze the conversion of inactive 11-dehydrocorticosterone to bioactive corticosterone, and its abnormally high expression in femoral head tissue is a key node in the occurrence and development of AIONFH **[46]**. The overactivation of this enzyme leads to an abnormal increase in the level of local active glucocorticoids in the femoral head. On the one hand, it induces the abnormal differentiation of bone marrow mesenchymal stem cells into adipocytes, resulting in the accumulation of bone marrow fat and compression of intraosseous capillaries, which destroys the microcirculation perfusion system of the femoral head and ultimately causes blood supply interruption. On the other hand, it directly inhibits the differentiation and activity of osteoblasts, and simultaneously promotes the activation and proliferation of osteoclasts, which breaks the physiological balance of bone metabolism, exacerbates the destruction of trabecular bone and osteocyte apoptosis, and drives the pathological progression of AIONFH from early ischemia to late structural collapse of the femoral head. Combined with the previous results of molecular docking and molecular dynamics simulation, it can be concluded that the core of the therapeutic effect of PNS on AIONFH lies in the stable binding of its active component G-Re to HSD11B1 and the targeted inhibition of its enzymatic activity. This effect can not only reduce the abnormal production of local active glucocorticoids in the femoral head and alleviate the mechanical compression of intraosseous blood vessels caused by the accumulation of bone marrow fat, but also synergistically activate angiogenic factors such as VEGF and FGF2 to effectively restore the blood supply in the ischemic area of the femoral head. Eventually, the vicious cycle of “AIONFH ischemia-osteonecrosis” is broken, and the targeted treatment of AIONFH is achieved through the multi-link synergistic effects of improving blood supply, regulating bone metabolism and inhibiting osteocyte apoptosis.

## 5 Innovations and Limitations of This Study

The innovation of this study lies in the construction of a complete research chain of in silico prediction-molecular validation-in vivo verification. Core targets were rapidly identified via network pharmacology combined with machine learning, which avoided the limitations of blind screening in traditional experiments. Molecular docking and molecular dynamics simulations were applied to elucidate the component-target interaction mechanism at the atomic level, providing microscopic evidence for the pharmacological effects of PNS. Animal experiments verified the therapeutic efficacy at both pathological and molecular levels, ensuring the reliability of the conclusions. Meanwhile, this study identified HSD11B1 as the key target of PNS in the treatment of AIONFH for the first time, which provides a new direction for the development of subsequent precision therapeutic targets.

However, the study still has certain limitations. The screening based on network pharmacology relied on existing databases, which may lead to the omission of low-abundance active components or unannotated targets. Molecular dynamics simulation was only performed on the single complex of HSD11B1 and Ginsenoside Re, without covering all core component-target combinations, thus failing to fully reflect the dynamic process of multi-target synergy. The animal model did not simulate the complications such as liver injury and metabolic syndrome that are often accompanied by human AIONFH. In addition, the intervention period was relatively short, and the effect of PNS on the long-term structural stability of the femoral head was not evaluated.

For future research, the CRISPR/Cas9 technology can be used to knock out core targets to clarify their necessity in the therapeutic effect of PNS. Monomer components of PNS can be isolated to evaluate their independent effects and analyze the specific mechanism of multi-component synergy. At the same time, a composite AIONFH model that is more consistent with clinical practice can be constructed to carry out long-term intervention experiments and small-sample clinical observations, so as to further verify the efficacy and safety of PNS.

## 6 Conclusion

This multi-dimensional study confirmed that the core mechanism of PNS in the treatment of AIONFH is the achievement of multi-link therapeutic effects of improving blood supply, inhibiting osteocyte apoptosis and balancing bone metabolism through the specific binding of its active components including Notoginsenoside R1 and Ginsenoside Rg1/Re to core targets such as FGF2, HSD11B1 and VEGFA, as well as the activation of signaling pathways including VEGF. Among them, the stable binding of Ginsenoside Re to HSD11B1 to inhibit glucocorticoid activation may represent the key molecular mechanism by which PNS improves the bone marrow microenvironment. This study not only provides solid experimental evidence for PNS as a potential therapeutic drug for AIONFH, but also proposes a research paradigm of combining computational biology with experimental medicine for the study on the multi-component and multi-target action mechanisms of traditional Chinese medicines. It is of great significance for promoting the modernization of traditional Chinese medicine and the development of novel therapeutic strategies for orthopedic diseases.

## List of abbreviations

(AIONFH): Alcohol-induced osteonecrosis of the femoral head
(PNS): Panax Notoginseng Saponins
(NG-R1): Notoginsenoside R1
(G-Rb1): Ginsenoside Rb1
(G-Rg1): Ginsenoside Rg1
(G-Re): Ginsenoside Re
(G-Rd): Ginsenoside Rd
(GEO): Gene Expression Omnibus
(HE): Hematoxylin-eosin
(TRAP): Tartrate-resistant acid phosphatase
(AKP): Alizarin Red and alkaline phosphatase
(RMSD): the Root Mean Square Deviation
(RMSF): Root Mean Square Fluctuation
(Rg): Radius of Gyration
(SASA): Solvent Accessible Surface Area
(qPCR): quantitative real-time polymerase chain reaction
(HSD11B1): 11β-hydroxysteroid dehydrogenase type 1
(VEGFA): vascular endothelial growth factor A
(FGF2): fibroblast growth factor 2
(MMP13): matrix metalloproteinase 13
(BMP2): bone morphogenetic protein 2
(KEGG): Kyoto Encyclopedia of Genes and Genomes
(BP): Biological Process
(CC): Cellular Component
(MF): Molecular Function
(SVM-RFE): upport vector machine-recursive feature elimination
(TYMS): thymidylate synthase
(Runx2): Runt-related transcription factor 2
(ANOVA): analysis of variance
(GB): Generalized Born

## Declarations

### Ethics approval and consent to participate

The Animal Experimental Ethics Committee Approval No. of Guangxi Zhuang Autonomous Region Academy of Chinese Medical Sciences was 2024093001. Clinical trial number: not applicable.

### Consent for publication

Not applicable

### Availability of data and materials

All data generated or analysed during this study are included in this published article [and its supplementary information files].

### Authors’ contributions

Bai R:wrote the main manuscript text

Shu H: prepared figures

Mo J:provided funding

Zhang X:reviewed the manuscript

Li Z:reviewed the manuscript

Chen X:reviewed the manuscript

Ye S:reviewed the manuscript

Nie X:reviewed the manuscript

Chen S:reviewed the manuscript

Liang B:provided funding

## Acknowledgements

This work was supported by the Guangxi Natural Science Foundation (Grant No. 2024GXNSFAA010370) and the Guangxi Traditional Chinese Medicine Appropriate Technology Development and Promotion Project (Grant No. GZSY2026008). The authors thank the supporting institutions and the Animal Experimental Ethics Committee of Guangxi Zhuang Autonomous Region Academy of Chinese Medical Sciences for ethical approval.

## References

[1] Chen, Wei et al. Zhongguo xiu fu chong jian wai ke za zhi = Zhongguo xiufu chongjian waike zazhi = Chinese journal of reparative and reconstructive surgery vol. 36, 11 (2022): 1420–1427. doi:10.7507/1002-1892.202206072

[2] Li, Zilin et al. “Advances in experimental models of osteonecrosis of the femoral head.” Journal of orthopaedic translation vol. 39 88–99. 3 Feb. 2023, doi:10.1016/j.jot.2023.01.003

[3] Kumar, Prasoon et al. “Association of Specific Genetic Polymorphisms with Atraumatic Osteonecrosis of the Femoral Head: A Narrative Review.” Indian journal of orthopaedics vol. 56, 5 771–784. 6 Jan. 2022, doi:10.1007/s43465-021-00583-3

[4] Lou, Yuhan et al. “Etiology, pathology, and treatment of osteonecrosis of the femoral head in adolescents: A comprehensive review.” Medicine vol. 103, 30 (2024): e39102. doi:10.1097/MD.0000000000039102

[5] Zhu, Weihong et al. “Bioengineering strategies targeting angiogenesis: Innovative solutions for osteonecrosis of the femoral head.” Journal of tissue engineering vol. 16 20417314241310541. 24 Jan. 2025, doi:10.1177/20417314241310541

[6] Chen, Xu et al. “Exploring the protective effects of PNS on acute myocardial ischaemia-induced heart failure by Transcriptome analysis.” Journal of ethnopharmacology vol. 271 (2021): 113823. doi:10.1016/j.jep.2021.113823

[7] Fan, Jing-Zheng et al. “Panax notoginseng saponins mitigate ovariectomy-induced bone loss and inhibit marrow adiposity in rats.” Menopause (New York, N.Y.) vol. 22, 12 (2015): 1343–50. doi:10.1097/GME.0000000000000471

[8] Yuan, Heng-Feng et al. “Protective effects of total saponins of panax notoginseng on steroid-induced avascular necrosis of the femoral head in vivo and in vitro.” Evidence-based complementary and alternative medicine : eCAM vol. 2015 (2015): 165679. doi:10.1155/2015/165679

[9] Yang, Wen-Jing et al. Zhongguo Zhong yao za zhi = Zhongguo zhongyao zazhi = China journal of Chinese materia medica vol. 48, 4 (2023): 1087–1097. doi:10.19540/j.cnki.cjcmm.20220926.501

[10] Feng, Guiyu et al. “Sequential Release of Panax Notoginseng Saponins and Osteopractic Total Flavone from Poly (L-Lactic Acid) Scaffold for Treating Glucocorticoid-Associated Osteonecrosis of Femoral Head.” Journal of functional biomaterials vol. 14, 1 31. 4 Jan. 2023, doi:10.3390/jfb14010031

[11] Qu, Jing et al. “Panax notoginseng saponins and their applications in nervous system disorders: a narrative review.” Annals of translational medicine vol. 8, 22 (2020): 1525. doi:10.21037/atm-20-6909

[12] Xiao, Haiyan et al. “Panax notoginseng saponins promotes angiogenesis after cerebral ischemia-reperfusion injury.” Journal of ginseng research vol. 48, 6 (2024): 592–602. doi:10.1016/j.jgr.2024.08.004

[13] Sun, Tao et al. “Effect of Panax notoginseng Saponins on Focal Cerebral Ischemia-Reperfusion in Rat Models: A Meta-Analysis.” Frontiers in pharmacology vol. 11 572304. 9 Feb. 2021, doi:10.3389/fphar.2020.572304

[14] Liu, Lu et al. “Effects of Panax notoginseng saponins on alleviating low shear induced endothelial inflammation and thrombosis via Piezo1 signalling.” Journal of ethnopharmacology vol. 335 (2024): 118639. doi:10.1016/j.jep.2024.118639

[15] Wei, Chen Chao et al. “Panax notoginseng saponins alleviate osteoporosis and joint destruction in rabbits with antigen-induced arthritis.” Experimental and therapeutic medicine vol. 22, 5 (2021): 1302. doi:10.3892/etm.2021.10737

[16] Zhang, Peng et al. “Network pharmacology: towards the artificial intelligence-based precision traditional Chinese medicine.” Briefings in bioinformatics vol. 25, 1 (2023): bbad518. doi:10.1093/bib/bbad518

[17] Li, Shao et al. Zhongguo Zhong yao za zhi = Zhongguo zhongyao zazhi = China journal of Chinese materia medica vol. 48, 22 (2023): 5965–5976. doi:10.19540/j.cnki.cjcmm.20230923.701

[18] Zhou, Meng-Nan et al. Zhongguo Zhong yao za zhi = Zhongguo zhongyao zazhi = China journal of Chinese materia medica vol. 46, 9 (2021): 2363–2369. doi:10.19540/j.cnki.cjcmm.20210222.201

[19] Yang, Xulong et al. “Treatment of liver fibrosis in hepatolenticular degeneration with traditional Chinese medicine: systematic review of meta-analysis, network pharmacology and molecular dynamics simulation.” Frontiers in medicine vol. 10 1193132. 11 May. 2023, doi:10.3389/fmed.2023.1193132

[20] Harrach, Michael F, and Barbara Drossel. “Structure and dynamics of TIP3P, TIP4P, and TIP5P water near smooth and atomistic walls of different hydroaffinity.” The Journal of chemical physics vol. 140, 17 (2014): 174501. doi:10.1063/1.4872239

[21] Yao, Chang-liang et al. “Simultaneous quantitation of five Panax notoginseng saponins by multi heart-cutting two-dimensional liquid chromatography: Method development and application to the quality control of eight Notoginseng containing Chinese patent medicines.” Journal of chromatography. A vol. 1402 (2015): 71–81. doi:10.1016/j.chroma.2015.05.015

[22] Yang, Xiaochen et al. “Protective effects of panax notoginseng saponins on cardiovascular diseases: a comprehensive overview of experimental studies.” Evidence-based complementary and alternative medicine : eCAM vol. 2014 (2014): 204840. doi:10.1155/2014/204840

[23] Zhou, Ping et al. “Ginsenoside Rb1 and mitochondria: A short review of the literature.” Molecular and cellular probes vol. 43 (2019): 1–5. doi:10.1016/j.mcp.2018.12.001

[24] Guo, Shasha et al. “Notoginsenoside R1: A systematic review of its pharmacological properties.” Die Pharmazie vol. 74, 11 (2019): 641–647. doi:10.1691/ph.2019.9534

[25] Yang, Chao et al. “Protein-Ligand Docking in the Machine-Learning Era.” Molecules (Basel, Switzerland) vol. 27, 14 4568. 18 Jul. 2022, doi:10.3390/molecules27144568

[26] Wu, Xiaodong et al. “Application of molecular dynamics simulation in biomedicine.” Chemical biology & drug design vol. 99, 5 (2022): 789–800. doi:10.1111/cbdd.14038

[27] Xia W, He L, Bao J, et al. Insights into small molecule inhibitor bindings to PD-L1 with residue-specific binding free energy calculation. J Biomol Struct Dyn. 2022;40(22):12277–12285.

[28] Coskun D, Chen W, Clark AJ, et al. Reliable and Accurate Prediction of Single-Residue pKa Values through Free Energy Perturbation Calculations. J Chem Theory Comput. 2022, 18(12):7193–7204.

[29] Zainal Ariffin, Shahrul Hisham et al. “Evaluation of in vitro osteoblast and osteoclast differentiation from stem cell: a systematic review of morphological assays and staining techniques.” PeerJ vol. 12 e17790. 25 Jul. 2024, doi:10.7717/peerj.17790

[30] Wick, Mark R. “The hematoxylin and eosin stain in anatomic pathology-An often-neglected focus of quality assurance in the laboratory.” Seminars in diagnostic pathology vol. 36, 5 (2019): 303–311. doi:10.1053/j.semdp.2019.06.003

[31] Jung, M, and R Kiesslich. “Chromoendoscopy and intravital staining techniques.” Bailliere’s best practice & research. Clinical gastroenterology vol. 13, 1 (1999): 11–9. doi:10.1053/bega.1999.0004

[32] Maguire, G A, and H Adnan. “An immunoprecipitation assay for high molecular weight alkaline phosphatase in human serum.” Annals of clinical biochemistry vol. 26 (Pt 2) (1989): 151–7. doi:10.1177/000456328902600211

[33] Komori T. Molecular Mechanism of Runx2-Dependent Bone Development. Mol Cells. 2020, 43(2):168–175.

[34] Zhou R, Huang R, Xu Y, Zhang D, Gu L, Su Y, Chen X, Shi W, Sun J, Gu P, Ni N, Bi X. Exosomes derived from mucoperiosteum Krt14+Ctsk+ cells promote bone regeneration by coupling enhanced osteogenesis and angiogenesis. Biomater Sci. 2024, 12(22):5753–5765.

[35] Gan, Di, and Changqing Zhang. Zhongguo xiu fu chong jian wai ke za zhi = Zhongguo xiufu chongjian waike zazhi = Chinese journal of reparative and reconstructive surgery vol. 27, 3 (2013): 365–8.

[36] Ahmed, Adeeba et al. “Induction of hepatic 11beta-hydroxysteroid dehydrogenase type 1 in patients with alcoholic liver disease.” Clinical endocrinology vol. 68, 6 (2008): 898–903. doi:10.1111/j.1365-2265.2007.03125.x

[37] Zhu, Kai et al. “Study on the mechanism of Shuanghe decoction against steroid-induced osteonecrosis of the femoral head: insights from network pharmacology, metabolomics, and gut microbiota.” Journal of orthopaedic surgery and research vol. 20, 1 202. 26 Feb. 2025, doi:10.1186/s13018-025-05619-0

[38] Requena-Ocaña, Nerea et al. “Vascular Endothelial Growth Factor as a Potential Biomarker of Neuroinflammation and Frontal Cognitive Impairment in Patients with Alcohol Use Disorder.” Biomedicines vol. 10,5 947. 20 Apr. 2022, doi:10.3390/biomedicines10050947

[39] Varoga, Deike et al. “Differential expression of vascular endothelial growth factor in glucocorticoid-related osteonecrosis of the femoral head.” Clinical orthopaedics and related research vol. 467, 12 (2009): 3273–82. doi:10.1007/s11999-009-1076-3

[40] Even-Chen, Oren et al. “FGF2 is an endogenous regulator of alcohol reward and consumption.” Addiction biology vol. 27, 2 (2022): e13115. doi:10.1111/adb.13115

[41] Huang, Gangyong et al. “FGF2 and FAM201A affect the development of osteonecrosis of the femoral head after femoral neck fracture.” Gene vol. 652 (2018): 39–47. doi:10.1016/j.gene.2018.01.090

[42] Prystupa, Andrzej et al. “Activity of MMP1 and MMP13 and amino acid metabolism in patients with alcoholic liver cirrhosis.” Medical science monitor : international medical journal of experimental and clinical research vol. 21 1008–14. 7 Apr. 2015, doi:10.12659/MSM.892312

[43] Chen, Gaoyang et al. “The expression of chondrogenesis-related and arthritis-related genes in human ONFH cartilage with different Ficat stages.” PeerJ vol. 7 e6306. 16 Jan. 2019, doi:10.7717/peerj.6306

[44] Gerjevic, Lisa Nicole et al. “Alcohol Activates TGF-Beta but Inhibits BMP Receptor-Mediated Smad Signaling and Smad4 Binding to Hepcidin Promoter in the Liver.” International journal of hepatology vol. 2012 (2012): 459278. doi:10.1155/2012/459278

[45] Kamiya, Nobuhiro et al. “Acute BMP2 upregulation following induction of ischemic osteonecrosis in immature femoral head.” Bone vol. 53, 1 (2013): 239–47. doi:10.1016/j.bone.2012.11.023

[46] Mukangwa M, Tetsuka M et al. Progesterone modulates HSD11B1-mediated cortisol production in luteinized bovine granulosa cells. J Reprod Dev. 2023;69(4):206–213. doi:10.1262/jrd.2023-005

